# Artefactual origin of biphasic cortical spike-LFP correlation

**DOI:** 10.1101/051029

**Authors:** Michael Okun

## Abstract

Electrophysiological data acquisition systems introduce various distortions into the signals they record. While such distortions were discussed previously, their effects are often not appreciated. Here I show that the biphasic shape of cortical spike-triggered LFP average (stLFP), reported in multiple studies, is likely an artefact introduced by high-pass filter of the neural data acquisition system when the actual stLFP has a single trough around the zero lag.

## Introduction

Cortical local field potential (LFP) is a readily measurable signal that provides a wealth of information on neuronal activity in the vicinity of the recording electrode. In particular, the relationship between LFP and the spiking of nearby neurons can lead to important insights into the cortical function, e.g., (Gray and Singer 1989; Arieli et al. 1995; Destexhe et al. 1999; Fries et al. 2001; Montemurro et al. 2008; Nauhaus et al. 2009; Martin and Schroeder 2016; Cui et al. 2016), and can be utilized in the design of brain-machine interfaces (BMIs) (Gulati et al. 2014; Hall et al. 2014). The simplest quantitative measure of the relationship between spiking activity and LFP is the spike-LFP cross-correlation, also known as spike-triggered LFP average (stLFP). stLFP can be computed for spike trains of individual neurons as well as for multi-unit spiking activity (MUA). In the former case, the shape and magnitude of stLFP are inherited from the cross-correlation between the membrane potential (Vm) of the neuron and the LFP (Okun et al. 2010).

In multiple previous studies, including our own, the stLFP often had a characteristic biphasic shape, whereby a trough around the 0 time lag is followed by a slower positivity, with a peak offset by several hundred milliseconds from 0, see Fig. 1 and (Arieli et al. 1995; Destexhe et al. 1999; Rasch et al. 2008; Rasch et al. 2009; Okun et al. 2010; Taub et al. 2015; Martin and Schroeder 2016). This positive peak of stLFP was thought to have a biophysical origin, and several possible explanations were proposed (Rasch et al. 2009; Ray 2014). Here, I show that in our data the positive peak of stLFP is a by-product of high-pass filtering of the actual LFP by the neural data acquisition system. Since passing the acquired signal through a high-pass filter with a cutoff frequency in the 0.1 – 1 Hz range is a standard feature of extracellular recording systems, it would only be natural to presume that the same explanation applies to other similar reports in the literature. Such interpretation also explains why in some studies, most notably when LFP was measured with an intracellular amplifier (i.e. effectively recording LFP in DC mode, which lets through all frequencies), a biphasic correlation between the LFP and the Vm of nearby neurons was not observed, e.g., (Poulet and Petersen 2008; Haider et al. 2016).

**Fig. 1.**
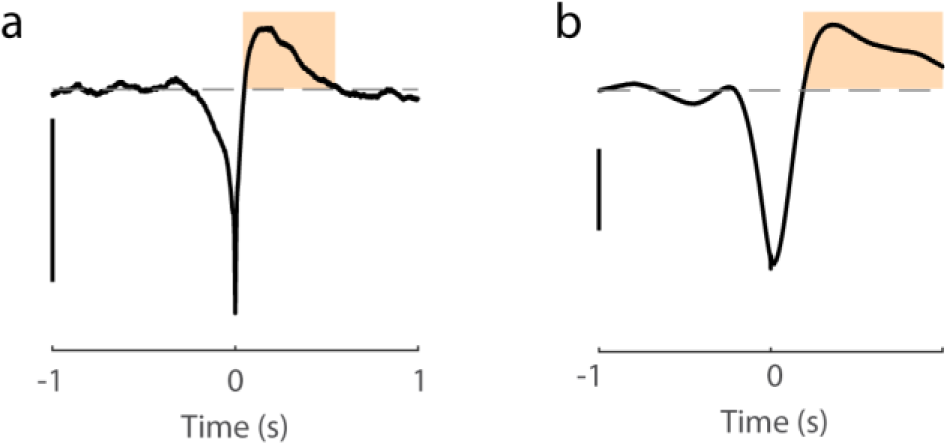
Examples of biphasic stLFP. **a** Recording performed with OpenEphys system. **b** Recording performed with Cerebus system. The positive peak which follows the central trough and produces the characteristic biphasic shape of stLFP is highlighted. Scale bars: 50 μV.

## Methods

All experimental procedures were conducted according to the UK Animals (Scientific Procedures) Act 1986 (Amendment Regulations 2012). Experiments were performed at University College London under personal and project licenses released by the Home Office following institutional ethics review.

The data analysed here originates from recordings of spontaneous activity made as part of previously published studies (Okun et al. 2015, 2016). Briefly, acute and chronic recordings were performed in the infragranular layers of primary visual cortex in C57BL/6J mice. In both cases recordings were performed in head fixed animals, after the mice underwent several sessions of acclimatisation to head fixation. Acute recordings were performed using Cerebus (Blackrock Microsystems, Salt Lake City, UT) neural acquisition system with Buzsaki32 probes (NeuroNexus, Ann Arbor, MI). The Cerebus system was used in the basic setting, where no digital filtering followed the initial high-pass at 0.3 Hz and low-pass at 7.5 kHz analogue filtering in the amplifier. The impedance of the recording sites was ˜1 MΩ at 1kHz. Chronic recordings were performed using the OpenEphys system (Siegle et al. 2015) and Intan RHD2132 16-channel amplifier board (Intan Technologies, Los Angeles, CA) with CM16 NeuroNexus probes with 2 tetrodes on each shank. The recordings were performed with the default filtering settings of OpenEphys software (1 Hz high-pass, 7.5 kHz low-pass). Prior to implantation, the probes were electroplated with the polymer PEDOT:PSS resulting in the recording sites’ impedance < 100 kΩ at 1kHz. For both acute and chronic recordings the signal was digitised at 30 kHz and stored for offline analysis. Spikes were detected using klusta software suite (Rossant et al. 2016). Here, only multiunit activity (MUA) comprised from all spikes detected on a shank or a tetrode was considered, hence spike sorting was not required. For stLFP computations, to reduce the contamination of the LFP with the spiking, I used MUA and LFP from different shanks of the probe, 150-200 μm apart.

Mathematically, high-pass filtering operation is a convolution of the original signal with the transfer function of the filter: *v_out_*(*t*) = *h* * *v*_*in*_(*t*), where *v_in_*(*t*) is the original signal, *v_out_*(*t*) is the recorded signal and h is the transfer function of the filter. According to the convolution theorem, in the frequency domain it holds that *V_out_*(*ω*) = *H*(*ω*)*V_in_*(*ω*), where capital letters denote the Fourier transforms of the original functions. *H*(*ω*) is a complex number, i.e., *H*(*ω*) = |H(*ω*)|*e*^*iargH*(*ω*)^, which is to say that in frequency *ω* the filter scales the amplitude of the original signal by |*H*(*ω*)| and shifts its phase by arg *H*(*ω*). In particular, for a high pass filter with cutoff at *ω*_0_, *H*(*ω*) ≈ 1 for frequencies *ω* >> *ω*_0_, and *H*(*ω*) ≈ 0 for *ω* ≪ *ω*_0_. Furthermore, if the transfer function *H* of the filter is known, it is possible to correct *v_out_*(*t*) offline, so that the phase shift introduced by the filter in frequencies around *ω*_0_ is undone, while the decrease in the power of these frequencies is retained. This is easily achieved by computing in the frequency domain a corrected signal *V_corr_*(*ω*) from *V_out_*(*ω*) in the following manner: *V_corr_*(*ω*) = *e^-iargH(ω)^V_out_(ω)*. A Matlab function that uses this approach for correcting OpenEphys recordings is now publicly available at https://github.com/open-ephvs/analvsis-tools/tree/master/lowFreqCorrection.

To examine the effect that high-pass filtering has on cross-correlation between signals, in addition to experimental spiking and LFP data, I used pairs of synthetic signals *x*(*t*), *y*(*t*) with pre-specified power spectrum *P*(*ω*) and coherence *C*(*ω*) (both are positive real valued functions, with *C*(*ω*) ≤ 1). *x*(*t*) and *y*(*t*) were constructed in the frequency domain in the following manner. *X*(*ω*) = *P*^1/2^(*ω*)*e*^*iφ*(*ω*)^, where for each frequency *ω*, *φ*(*ω*) was drawn randomly and uniformly from the [0,2π) interval. *Y*(*ω*) = −*C*(*ω*)*X*(*ω*) + *Y*_2_(*ω*), where *Y*_2_(*ω*) is an additional signal with a power spectral density of (1 – *C*^2^(*ω*))*P*(*ω*). *Y*_2_(*ω*) was generated similarly to *X*(*ω*), however the phases of *X*(*ω*) and *Y*_2_(*ω*) were drawn independently, thus the two signals were uncorrelated (and incoherent) with each other. Finally, the time domain signals *x*(*t*), *y*(*t*) were obtained by taking the inverse Fourier transform of *X*(*ω*) and *Y*(*ω*). The result of applying a high-pass filter *h* to *y*(*t*) was also computed in the frequency domain, by taking the inverse Fourier transform of *H*(*ω*)*Y*(*ω*).

## Results

The way low frequency signals are processed by the neural data acquisition system is central to the understanding of the stLFP waveform. To measure the transfer function of an acquisition system, I connected a function generator (TG310, Thurlby Thandar Instruments, UK) to the head-stage of the amplifier, and delivered sine waves of equal amplitude (˜5 mV) spanning frequencies between 30 Hz and 0.03 Hz. The signal generator also emitted a TTL signal, which marked the extrema of the input sine wave. Comparing the TTL signal to the output of the amplifier allowed estimating the phase shift introduced by its filters, while comparing the amplitude of the output across frequencies provided an estimate of the gain (Fig. 2a, b). In this manner I obtained the transfer function for Cerebus and OpenEphys systems. The Cerebus system was tested in the basic setting (also used for recordings), where no digital filtering followed the initial analog filtering in the amplifier. The OpenEphys system was tested with 1 Hz cutoff frequency of the high-pass filter, which is the default of its data acquisition software, and with 0.1 Hz cutoff, which is the lowest possible value in this system. The results of these measurements show that amplifier filtering substantially shifts phases even in frequencies > 1 Hz (Fig. 2c, d). Surprisingly, the phase shift was larger for OpenEphys with a nominal 0.1 Hz cutoff frequency than for Cerebus with a nominal 0.3 Hz cutoff frequency (Fig. 2d), demonstrating that the cutoff frequency on its own (without a full specification of the type and order of the filter) does not provide a complete characterisation of the highpass filtering properties of the amplifier.

**Fig. 2.**
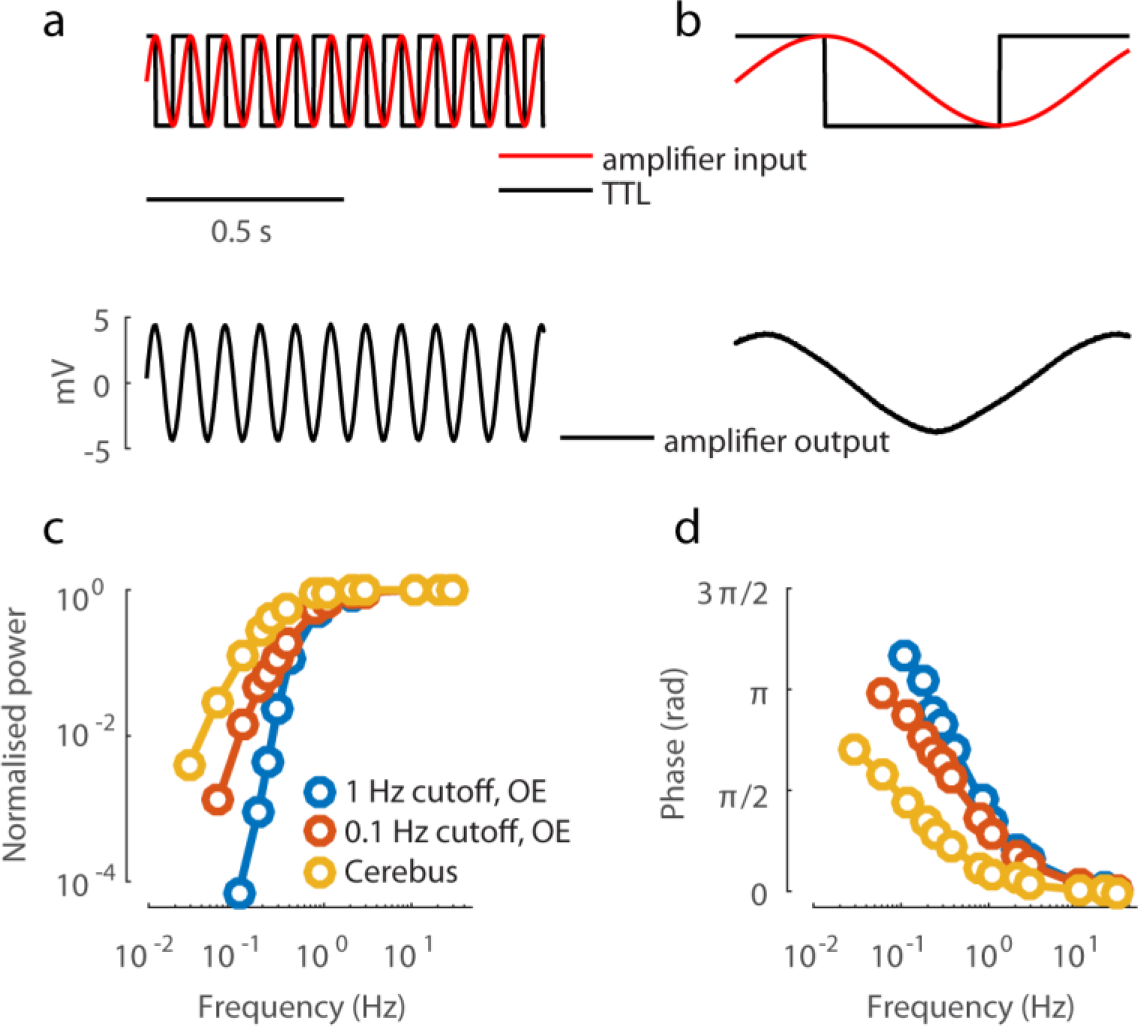
Amplifier transfer function. **a** Measuring the amplifier transfer function. Output of the function generator (red sine wave) was fed into the head-stage of the amplifier, and the TTL signal (overlaid, black) was stored for offline analysis. The output of the amplifier (black sine wave) was also stored for offline analysis. **b** As in a, for a sinusoidal input of lower frequency. A clear phase shift between the input (red) and output (black) sine waves is seen, with the output leading the input. A small reduction in amplitude (cf. a) is also apparent. **c, d** The gain and the phase shift introduced by the amplifier, measured across a range of frequencies as demonstrated in a-b, for OpenEphys (OE) system with 1 Hz and 0.1 Hz cutoffs, and for the Cerebus system.

To understand the effect such high-pass filtering has on the LFP and its correlation with spikes or Vm of nearby neurons, I started by considering pairs of synthetic signals *x*(*t*), *y*(*t*) constructed (see Methods) to have a symmetric correlation on the same timescale as empirical stLFPs (Fig. 3a, b). In this analysis, *x*(*t*)represents spikes or membrane potential, while *y*(*t*) represents the LFP. After the transfer functions of the amplifiers were applied to *y*(*t*), its correlation with *x*(*t*) became biphasic, similar to the observed cortical stLFP (Fig. 3c, cf. Fig. 1). Next, I examined the effect that the power spectrum of *y*(*t*) has on the shape of the cross-correlation with *x*(*t*) before and after the distortion by high-pass filtering. When *y*(*t*)contains more power in frequencies < 10 Hz, its correlation with *x*(*t*) increases in magnitude (Fig. 3d, e). In these cases, the distortion introduced by high-pass filtering of *y*(*t*) is much more prominent and is manifested in two features of the *x*(*t*)-*y*(*t*) cross-correlation. First, the artefactual positive peak can grow to be of almost equal magnitude to the central negative trough around 0 time lag (Fig. 3f). Second, the correlation at 0 time lag differs prominently from its true value (cf. Fig. 3e, f).

**Fig. 3.**
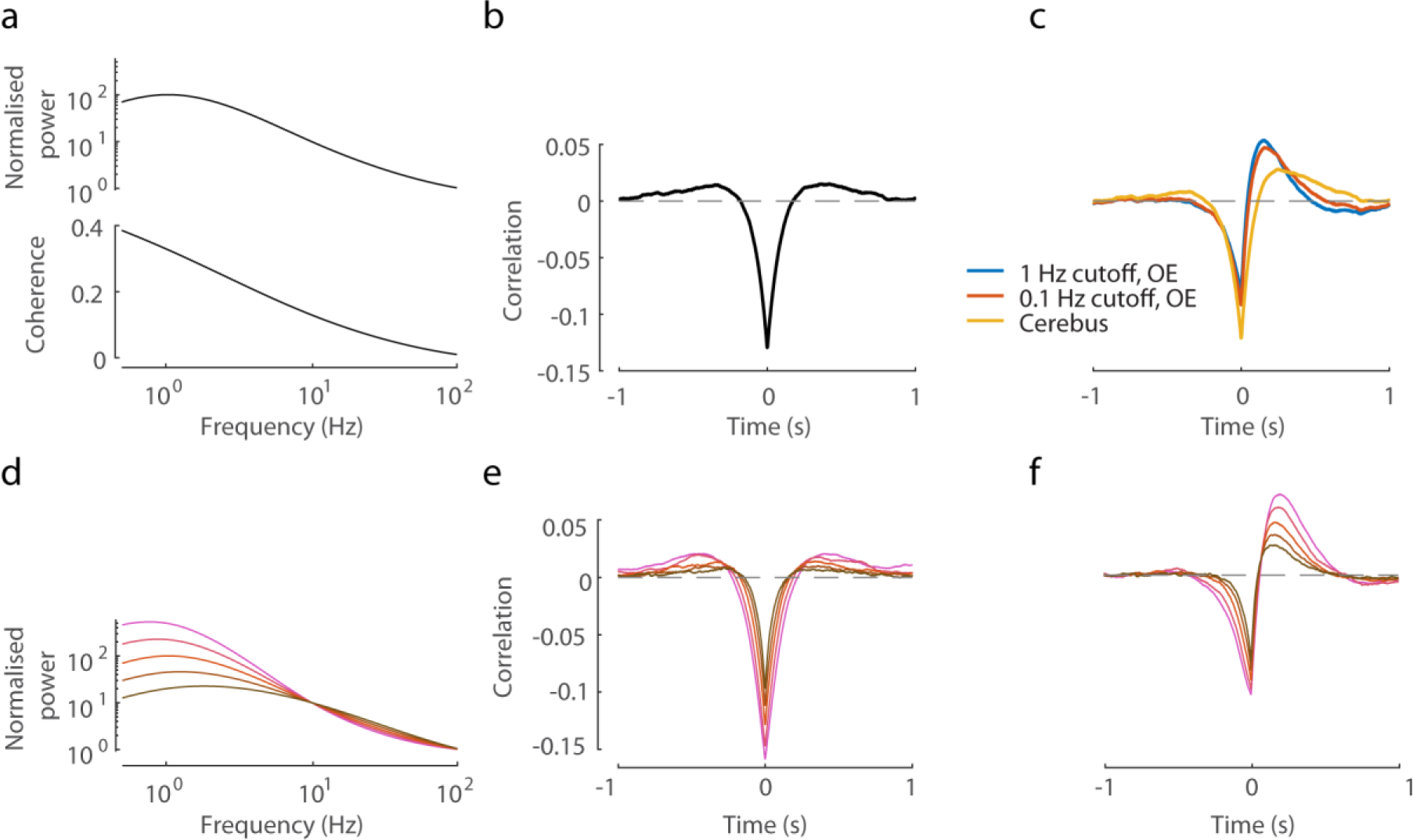
Simulation of the effect of amplifier high-pass filter on measured cross-correlations. **a** The power spectrum and coherence of a synthetically generated pair of signals *x*(*t*) and *y*(*t*) (see Methods). **b** The symmetric crosscorrelation between *x*(*t*) and *y*(*t*). **c** The cross-correlation after *y*(*t*) was filtered with the amplifier transfer functions shown in Fig. 2c-d. **d** Pairs of synthetic signals *x*(*t*), *y*(*t*) were generated where the power spectrum of *x*(*t*) and the coherence were as in a, while the power spectrum of *y*(*t*) varied as shown (the middle spectrum is as in a). **e** The cross-correlation between *x*(*t*) and *y*(*t*), for the different cases of *y*(*t*)’s power spectrum shown in d (colours match). **f** The cross-correlations after *y*(*t*) was filtered with the 0.1 Hz cutoff OpenEphys high-pass transfer function (colours match to d-e).

When the above simulations were repeated using a spike train derived from *x*(*t*) instead of the continuous signal *x*(*t*) itself (spikes corresponded to *x*(*t*) going above its 99th percentile value), the results were identical up to a scaling factor of the ordinate (data not shown). This result is consistent with the mathematical theory of triggered cross-correlations (Boer and Kuyper 1968).

The knowledge of the transfer function of the amplifier allows to reverse, via offline processing, the phase distortion the amplifier introduces into the LFP recording, as described in the Methods section. Here I applied such a correction to recordings previously acquired in the mouse primary visual cortex. Examples of LFP traces before and after the phase correction are shown in Fig. 4a, b. As expected, the difference between the recorded and the corrected signals is in low frequencies, and at first glance might appear rather small. However, after the phase correction, the correlation with the spiking activity was no longer biphasic (Fig. 4c, d).

**Fig. 4.**
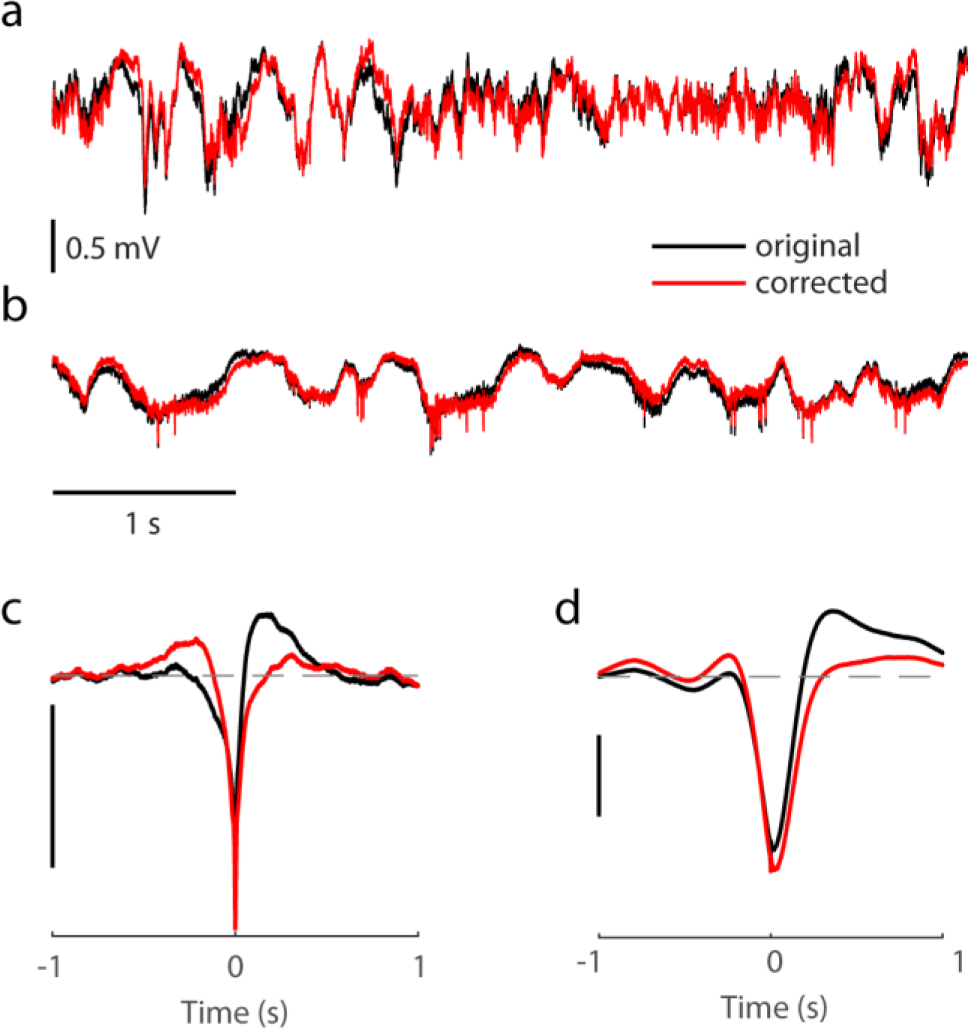
Offline correction of phase distortion. **a** Example of LFP in mouse primary visual cortex, recorded with the OpenEphys system with 1 Hz cutoff high-pass filter, before and after the phase correction. **b** As in a, for LFP exhibiting a rather different dynamics, recording performed with the Cerebus system. **c, d** stLFP for recordings shown in a, b, computed using LFP signal as it was recorded by the data acquisition system and after phase correction. Scale bars: 50 μV.

## Discussion

A comprehensive overview of the distortions caused by recording electrodes and amplifier filtering was provided in (Nelson et al. 2008). Here I did not consider distortions introduced by the silicon probe because according to their measurements such a distortion is relatively small (< 0.33 rad) for amplifiers with input impedance > 1 GΩ (which, according to the specifications, is the case for both Cerebus and Intan amplifier head-stages). The methodological approach to measuring and correcting the distortion used here is virtually identical to the approach proposed in (Nelson et al. 2008). However, Nelson et al. do not provide any concrete examples, beyond a passing mention of numerous publications where LFP distortion might not have been accounted for. The present work therefore is, to the best of my knowledge, the first to provide a concrete example of a well-documented feature of spike-LFP dynamics that appears to be produced by amplifier filtering rather than genuine biophysical mechanisms. That being said, this explanation does not fully exclude the possibility that under some experimental conditions cortical stLFP can have a biphasic shape. More generally, the present work demonstrates that low frequency filtering, which is employed by the vast majority of neural data acquisition systems in use today, has important confounding implications for the study of LFP. In particular, the distortion is not limited to stLFP, but is a general property of cross-correlations involving the LFP signal. In other words, the second signal can be neural or external events other thank spikes, e.g., onsets of UP states (Lewis et al. 2012).

The distortion of the low frequencies of the LFP can be corrected offline. This requires measuring the transfer function of the amplifier system and reversing its effect, as demonstrated here and in (Nelson et al. 2008). An example Matlab code to perform this correction is provided in supplementary material of (Nelson et al. 2008), and some vendors might already have a special utility for their neural data acquisition systems (e.g., FPAlign of Plexon Inc, Dallas, TX). A Matlab function for correcting OpenEphys recordings (which was used to generate Fig. 4a, c) is publicly available at https://github.com/open-ephys/analysis-tools/tree/master/lowFreqCorrection.

## Acknowledgements

I would like to thank Sylvia Schroeder and Nicholas Steinmetz for commenting on the manuscript and Kenneth Harris and Matteo Carandini for supporting this work (via Wellcome Trust grants 95668 and 95669).

## References

Arieli, A., Shoham, D., Hildesheim, R., & Grinvald, A. (1995). Coherent spatiotemporal patterns of ongoing activity revealed by real-time optical imaging coupled with single-unit recording in the cat visual cortex. Journal of Neurophysiology, 73(5), 2072–2093.

Boer, E. D., & Kuyper, P. (1968). Triggered Correlation. IEEE Transactions on Biomedical Engineering, BME-15(3), 169–179. doi:10.1109/TBME.1968.4502561

Cui, Y., Liu, L. D., McFarland, J. M., Pack, C. C., & Butts, D. A. (2016). Inferring Cortical Variability from Local Field Potentials. The Journal of Neuroscience, 36(14), 4121–4135. doi:10.1523/JNEUROSCI.2502-15.2016

Destexhe, A., Contreras, D., & Steriade, M. (1999). Spatiotemporal analysis of local field potentials and unit discharges in cat cerebral cortex during natural wake and sleep states. The Journal of Neuroscience, 19(11), 4595–4608.

Fries, P., Reynolds, J. H., Rorie, A. E., & Desimone, R. (2001). Modulation of oscillatory neuronal synchronization by selective visual attention. Science (New York, N.Y.), 291(5508), 1560–1563. doi:10.1126/science.291.5508.1560

Gray, C. M., & Singer, W. (1989). Stimulus-specific neuronal oscillations in orientation columns of cat visual cortex. Proceedings of the National Academy of Sciences of the United States of America, 86(5), 1698–1702.

Gulati, T., Ramanathan, D. S., Wong, C. C., & Ganguly, K. (2014). Reactivation of emergent task-related ensembles during slow-wave sleep after neuroprosthetic learning. Nature Neuroscience, 17(8), 1107–1113. doi:10.1038/nn.3759

Haider, B., Schulz, D. P. A., Häusser, M., & Carandini, M. (2016). Millisecond Coupling of Local Field Potentials to Synaptic Currents in the Awake Visual Cortex. Neuron, 90(1), 35–42. doi:10.1016/j.neuron.2016.02.034

Hall, T. M., Nazarpour, K., & Jackson, A. (2014). Real-time estimation and biofeedback of single-neuron firing rates using local field potentials. Nature Communications, 5, 5462. doi:10.1038/ncomms6462

Lewis, L. D., Weiner, V. S., Mukamel, E. A., Donoghue, J. A., Eskandar, E. N., Madsen, J. R., et al. (2012). Rapid fragmentation of neuronal networks at the onset of propofol-induced unconsciousness. Proceedings of the National Academy of Sciences, 109(49), E3377–E3386. doi:10.1073/pnas.1210907109

Martin, K. A. C., & Schroeder, S. (2016). Phase Locking of Multiple Single Neurons to the Local Field Potential in Cat V1. The Journal of Neuroscience, 36(8), 2494–2502. doi:10.1523/JNEUROSCI.2547-14.2016

Montemurro, M. A., Rasch, M. J., Murayama, Y., Logothetis, N. K., & Panzeri, S. (2008). Phase-of-Firing Coding of Natural Visual Stimuli in Primary Visual Cortex. Current Biology, 18(5), 375–380. doi:10.1016/j.cub.2008.02.023

Nauhaus, I., Busse, L., Carandini, M., & Ringach, D. L. (2009). Stimulus contrast modulates functional connectivity in visual cortex. Nature neuroscience, 12(1), 70–76. doi:10.1038/nn.2232

Nelson, M. J., Pouget, P., Nilsen, E. A., Patten, C. D., & Schall, J. D. (2008). Review of signal distortion through metal microelectrode recording circuits and filters. Journal of Neuroscience Methods, 169(1), 141–157. doi:10.1016/j.jneumeth.2007.12.010

Okun, M., Lak, A., Carandini, M., & Harris, K. D. (2016). Long Term Recordings with Immobile Silicon Probes in the Mouse Cortex. PLOS ONE, 11(3), e0151180. doi:10.1371/journal.pone.0151180

Okun, M., Naim, A., & Lampl, I. (2010). The subthreshold relation between cortical local field potential and neuronal firing unveiled by intracellular recordings in awake rats. The Journal of Neuroscience, 30(12), 4440–4448. doi:10.1523/JNEUR0SCI.5062-09.2010

Okun, M., Steinmetz, N. A., Cossell, L., lacaruso, M. F., Ko, H., Barthó, P., et al. (2015). Diverse coupling of neurons to populations in sensory cortex. Nature, 521(7553), 511–515. doi:10.1038/nature14273

Poulet, J. F. A., & Petersen, C. C. H. (2008). Internal brain state regulates membrane potential synchrony in barrel cortex of behaving mice. Nature, 454(7206), 881–885. doi:10.1038/nature07150

Rasch, M. J., Gretton, A., Murayama, Y., Maass, W., & Logothetis, N. K. (2008). Inferring spike trains from local field potentials. Journal of neurophysiology, 99(3), 1461–1476. doi:10.1152/jn.00919.2007

Rasch, M., Logothetis, N. K., & Kreiman, G. (2009). From neurons to circuits: linear estimation of local field potentials. The Journal of neuroscience, 29(44), 13785–13796. doi:10.1523/JNEUROSCI.2390-09.2009

Ray, S. (2014). Challenges in the quantification and interpretation of spike-LFP relationships. Current Opinion in Neurobiology, 31C, 111–118. doi:10.1016/j.conb.2014.09.004

Rossant, C., Kadir, S. N., Goodman, D. F. M., Schulman, J., Hunter, M. L. D., Saleem, A. B., et al. (2016). Spike sorting for large, dense electrode arrays. Nature Neuroscience, 19(4), 634–641. doi:10.1038/nn.4268

Siegle, J. H., Hale, G. J., Newman, J. P., & Voigts, J. (2015). Neural ensemble communities: open-source approaches to hardware for large-scale electrophysiology. Current Opinion in Neurobiology, 32, 53–59. doi:10.1016/j.conb.2014.11.004

Taub, A. H., Lampl, I., & Okun, M. (2015). Local Field Potential, Relationship to Membrane Synaptic Potentials. In D. Jaeger & R. Jung (Eds.), Encyclopedia of Computational Neuroscience (pp. 1572–1579). Springer New York. http://link.springer.com/referenceworkentry/10.1007/978-1-4614-6675-8_728.

